# Allergic inflammation hinders synergistic viral-bacterial co-infection in C57BL/6 mice

**DOI:** 10.1101/550459

**Authors:** Kim S. LeMessurier, Amy R. Iverson, Ti-Cheng Chang, Maneesha Palipane, Peter Voge, Jason W. Rosch, Amali E. Samarasinghe

## Abstract

Asthma is a chronic airways disease that can be exacerbated during respiratory infections. Our previous findings that the inflammatory state of allergic airways at the time of influenza A virus (IAV) infection in combination with epidemiologic findings that asthmatics were less likely to suffer from severe influenza during the 2009 pandemic suggest that additional complications of influenza, such as increased susceptibility to bacterial superinfection, may be mitigated in the allergic host. To test this hypothesis, we developed a murine model of ‘triple-disease’ in which mice were first rendered allergic to *Aspergillus fumigatus* and co-infected with IAV and *Streptococcus pneumoniae* seven days apart. Significant alterations to known synergistic effects of co-infection were noted in the allergic mice including reduced morbidity and mortality, bacterial burden, maintenance of alveolar macrophages, and reduced lung inflammation and damage. The lung microbiome of allergic mice differed from that of non-allergic mice during co-infection. To investigate the impact of the microbiome on the pathogenesis of lung disease, we induced a perturbation with a short course of fluoroquinolone antibiotic that is often prescribed for lung infections. A significant change in the microbiome was complemented with alterations to the inflammatory profile and a drastic increase in pro-inflammatory cytokines in allergic mice which were now susceptible to severe disease from IAV and *S. pneumoniae* co-infection. Our data suggest that responses to co-infection in allergic hosts likely depends on the immune and microbiome states and that antibiotics should be used with caution in individuals with underlying chronic lung disease.

**Author Summary**

Asthma is a condition of the lungs that affects millions worldwide. Traditionally, respiratory infections are considered to have a negative impact on asthmatics. However, epidemiological data surrounding the 2009 influenza pandemic suggest that asthmatics may be better equipped to counter severe influenza including bacterial pneumonia. Herein, we introduce a novel mouse model system designed to recapitulate an influenza virus and Streptococcal co-infection in a host with fungal asthma. We found that underlying allergic asthma protects against severe disease induced by co-infection. Mice with underlying allergic inflammation had reduced damage to the lungs and did not show signs of respiratory distress. Among the differences noted in the allergic mice that were protected from viral and bacterial co-infection, was the lung microbiome. Allergic mice lost their protection from co-infection after we perturbed their lung microbiome with antibiotics suggesting that the lung microbiome plays a role in host immunity against invading pathogens.

## Introduction

Lung diseases are a leading cause of morbidity and mortality worldwide. Acute and chronic respiratory diseases, excluding infections, affect greater than 12% of the population in the United States and hundreds of millions worldwide (1). Asthma is the most prevalent of these (2) and has the greatest economic burden (3), in addition to being one of most challenging lung conditions to investigate and treat. While the exact etiology of asthma remains unstipulated, specified phenotypes based on symptoms, immunologic profiles, genes, and environment, are confounded by gender and age. Furthermore, asthma exacerbations can be triggered by respiratory viral infections (4, 5) and some reports suggest that asthmatics are at risk for bacterial pneumonia (6, 7).

Over four million deaths every year (predominantly children <5 years) are caused by acute respiratory infections (8). Influenza and Streptococcal pneumonia contribute to approximately one million hospitalizations annually in the U.S. (9, 10). While influenza alone can be fatal, recovering patients have an increased susceptibility to respiratory bacterial infections (11), of which *Streptococcus pneumoniae* (*Spn*), a pathogen associated with community-acquired pneumonia (12), is highly associated with severe disease during influenza infections. The ‘Spanish Flu’ pandemic exemplified this predilection with the majority of deaths attributed to subsequent bacterial infections (13), as did the 2009 Swine Flu pandemic in which 29-55% of deaths resulted from secondary bacterial pneumonia (14, 15). The occurrence of pulmonary infections in asthmatics is augmented by high disease incidences of each, and seasonal overlap between infectious agents and allergens.

Although asthma was a risk factor for hospitalization during the 2009 ‘Swine Flu’ pandemic (16). subsequent studies noted that some asthmatics had less severe outcome (including reduced bacterial pneumonia) compared to non-asthmatics (17-21). Explanations for this unexpected and counterintuitive outcome are sparse, although possibilities include steroid use (22), heightened medical care, and increased likelihood of vaccinations in asthmatics (21). A better understanding of mechanisms that promote protection from pulmonary infection-mediated complications in asthmatics is important and necessary, but currently lacking in the literature.

Effective interrogation of host-pathogen interactions during allergic asthma and respiratory infections necessitates a single animal model that can capture the nuances of both morbidities. Our mouse model of fungal asthma and influenza (23) effectively recapitulates epidemiologic findings that some asthmatics had less severe disease during the 2009 influenza A virus (IAV) pandemic (20, 24). To investigate viral-bacterial synergy in the allergic host, we developed a ‘triple-disease model’ by combining our fungal asthma model (25) with a well-established model of IAV and *Spn* co-infection (26). Our findings suggest that pre-existing allergic asthma protects the host from severe morbidity, as shown by maintenance of weight, and reduced viral-bacterial synergism. Allergic mice also had reduced bacterial burdens, altered inflammatory cell profiles (more eosinophils and macrophages and fewer neutrophils) as well as a distinct lung microbiome compared to those with IAV and *Spn* co-infection alone. Inducing dysbiosis with antibiotics caused a partial reversal of this protective phenotype observed in the allergic mice.

## Results

Asthmatics are generally considered to have heightened susceptibility to respiratory infections (27). However, exact reasons for this outcome are not well established in humans due to inaccessibility of mucosal tissue samples limiting thorough mechanistic interrogations. Therefore, animal model systems are invaluable in such instances, and yet, those that allow the interrogation into the convergence of immunologically distinct conditions like asthma, influenza, and pneumococcal pneumonia are lacking. Recent work with our animal model of asthma and influenza comorbidity underscored the importance of the state of the allergic airways at the time of viral infection in the pathogenesis of influenza in allergic hosts (23), and identified a novel antiviral function for eosinophils (28). Since viral-bacterial synergy has previously established to cause severe pneumonia and mortality (29), we sought to determine the effect of an allergic microenvironment in the lung on subsequent co-infections with IAV and *Spn*, both alone and in combination.

### Allergic airways inflammation protected mice against severe disease from co-infection

Mouse model systems that can simulate complex interactions between allergy and respiratory infections are limited, but important to study disease-disease interactions that may alter host responses. Since respiratory infections with viruses and bacteria are considered triggers for the development of asthma, the previously reported animal models utilized infectious agents prior to allergen provocation (30). While asthma can indeed be triggered by respiratory infections, it can also be exacerbated by the same (31, 32). Herein, our first goal was to develop and characterize a model system in which respiratory infections occurred in established allergic airways disease. Mice were subjected to *A. fumigatus* allergen sensitization and challenge (25, 33), and then infected with IAV (23), followed by *S. pneumoniae* (**Fig 1A**). Naïve mice were used to measure baseline, while asthma-only, influenza-only (Flu Ctr), bacteria-only (Bact Ctr) mice served as single disease controls. Dual condition groups included Asthma+Flu (AF), Asthma+Bact (AB), and Flu+Bact (FB), while Asthma+Flu+Bact (AFB) triple-disease condition served as the experimental group.

**Figure 1.**
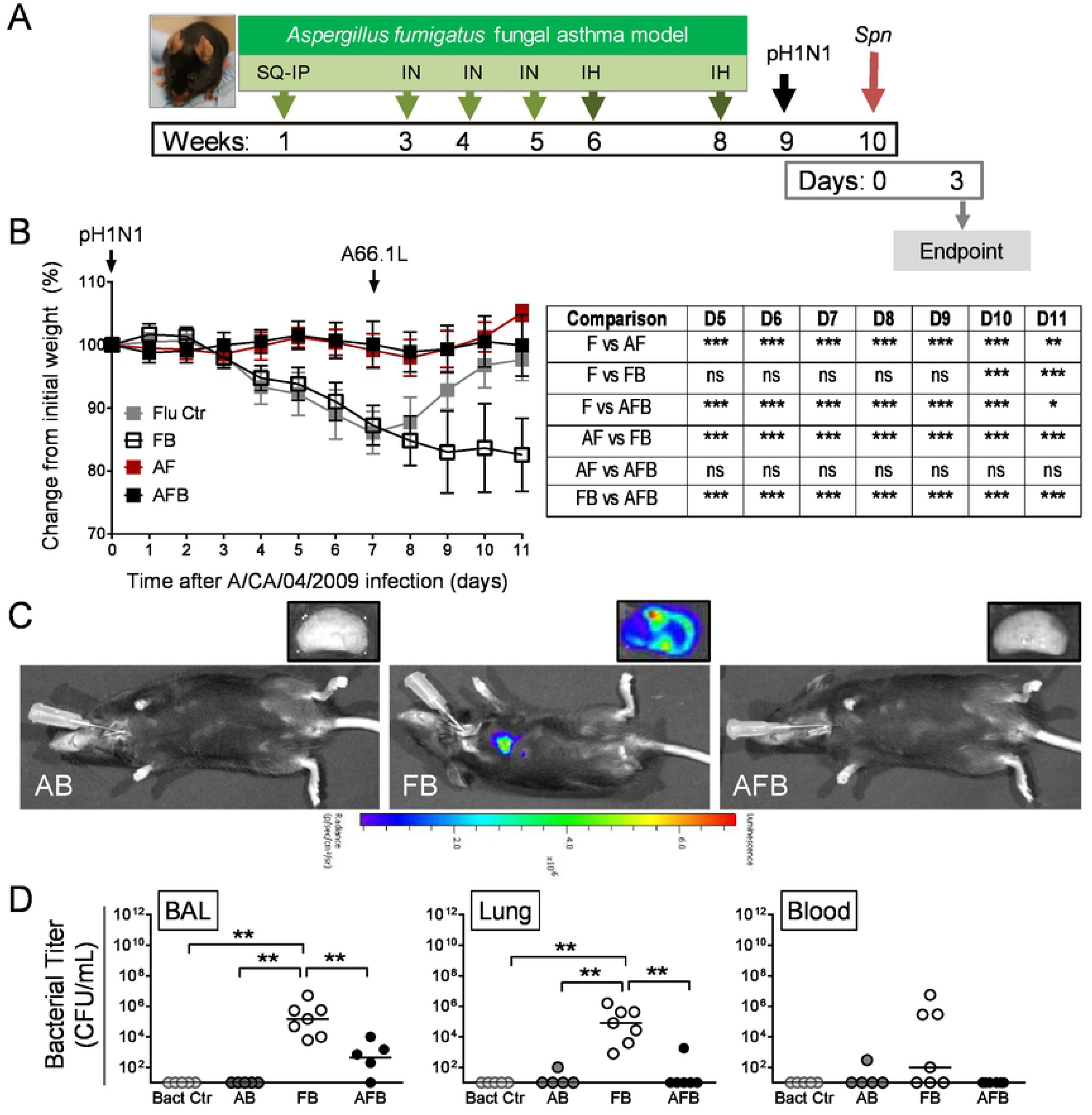
Allergic airways disease protects the host from synergistic morbidity from influenza and bacterial pneumonia. Timeline of triple-disease model (A) wherein allergen sensitized and challenged mice are infected with influenza A virus (pH1N1) and *Streptococcus pneumonia* (*Spn*). Weight loss in each group and comparative statistics associated with weight loss (B). Bioluminescence reading for bacteria in mice and harvested lung lobes (C). Bacterial burden in the bronchoalveolar lavage (BAL), lung homogenate, and blood (D). Data are representative of one study from four independent studies. n=5-7 mice per group. Data analysed by one-or two-way ANOVA and *P<0.05, **P<0.01, and ***P<0.001. SQ: subcutaneous; IN: intranasal; IH: inhalation; F and Flu: influenza; B and Bact: bacteria; A: asthma.

As previously demonstrated by us (23), IAV infection during peak airways inflammation did not induce weight loss in allergic mice (AF group, **Fig 1B**), whereas the same dose of virus triggered about a 12% weight loss in non-allergic mice (F group, **Fig 1B**). Non-allergic mice that were co-infected (FB group) lost ∼20% weight at the termination point in this study (**Fig 1B**), and continued to lose weight and succumbed to disease by 6 dpi with *Spn* (data not shown). In stark contrast, allergic mice that were subsequently co-infected (AFB group) did not lose weight and had a comparable weight profile to the AF group (**Fig 1B**) and 83% in the AFB group survived (data not shown). As such, although mortality was not an output of this study, allergy appeared to delay/protect mice from IAV+*Spn*-induced mortality which may provide a time advantage for clinical therapeutics. Infectious virus was absent in the lungs of mice in all groups at 3 dpi (data not shown) which differs from previous studies that have demonstrated a viral rebound after *Spn* co-infection, albeit using the laboratory strain of IAV (34). The bacterial burden in the allergic lungs was not sufficient to visualize by fluorescence like in co-infection alone (**Fig 1C**), but conventional enumeration of pneumococci on blood agar showed significant reduced bacterial loads in allergic mice compared to the non-allergic co-infected mice (**Fig 1D**). Bacterial dissemination into the blood may also be delayed/reduced in allergic mice (**Fig 1D**).

### Allergic mice had a more diverse immune cell signature in the airways although tissue inflammation was lower compared to non-allergic mice during co-infection

Inflammation is an important hallmark of both allergic disease and respiratory infections, although the cell types that dominate are different. We measured the number and type of leukocytes in the airways (bronchoalveolar lavage, BAL) as a marker of disease severity. As expected, inflammatory cells were increased significantly over steady state (3.70×10^4^± 1.04×10^4^live cells) after each trigger (**Fig 2A**). Macrophages were more abundant in single infections with and without allergy, but significantly lower in the FB group (**Fig 2B**) as previously demonstrated (35). Eosinophils, B cells, and CD4^+^T cells followed a similar pattern of abundance, wherein cell numbers were significantly higher in pathogen infected allergic mice than their non-allergic counterparts (**Fig 2B**). Neutrophil infiltration was markedly higher in the FB group, while CD8^+^T cell recruitment was similar except for the asthma-and bacteria-only controls which had very low influx into airways (**Fig 2B**).

**Figure 2.**
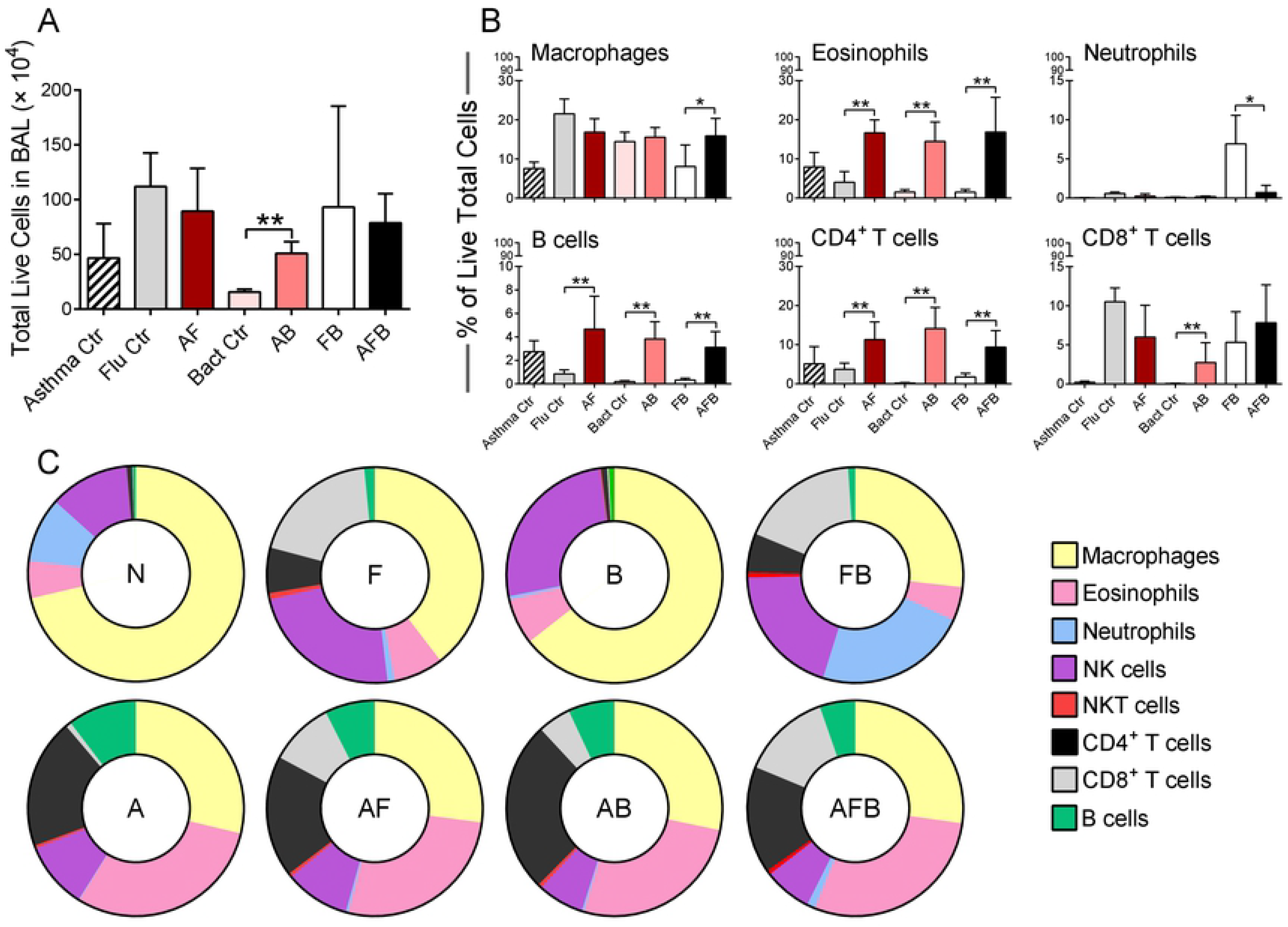
Inflammatory cell profile in the bronchoalveolar lavage compartment of mice after triple-disease model. Cells purified from the BAL were enumerated (A) and used for flow cytometry to identify changes in cell types (B). The percentage of each cell type was used to normalize data to identify differences in these cell types in the airways during various disease conditions (C). Data are representative of one study from four independent studies. n=5-7 mice per group. Data were analysed by Mann-Whitney test against the direct control group for each experimental condition. *P<0.05 and **P<0.01. Ctr: control; F and Flu: influenza virus; B and Bact: bacteria; A: asthma; N: naïve.

Over exuberant immune responses are considered a mechanism by which synergistic actions of IAV and *Spn* increased host morbidity and mortality (36). Analysis of hematoxylin and eosin stained sections of lung tissue showed that widespread parenchymal inflammation was present in IAV-infected mice (Flu, **Fig 3**) but that *Spn*-infected mice had minimal areas of inflammation (Bacteria, **Fig 3**) most likely due to effective bacterial clearance in otherwise healthy hosts. In contrast, extensive areas of lung parenchyma in FB group mice were consolidated by fluid exudates and inflammatory cells consisting mostly of neutrophils and macrophages (**Fig 3**). As expected, inflammation in the asthma-only control mice mostly surrounded the terminal airways (**Fig 3**), and similar inflammatory foci were observed around the small airways in both the AF (**Fig 3**) and AB groups (**Fig 3**). Significantly, pulmonary lesions in the AFB group mice (**Fig 3**) were much less severe than those in the FB group (**Fig 3**), and resembled those of AF mice (**Fig 3**). Histopathologic scoring of diffuse alveolar damage markers such as alveolar inflammation and protein/fibrin deposition were all much higher in the FB group than in the AFB. The higher levels of mucus production in all allergic mice, irrespective of the presence or type of infectious agent, correlated with the reduced damage and loss of bronchiolar epithelium in these lungs. These analyses suggest that IAV infection may provide the dominant antigen triggers for the resulting and subsequent inflammation in the lungs.

**Figure 3.**
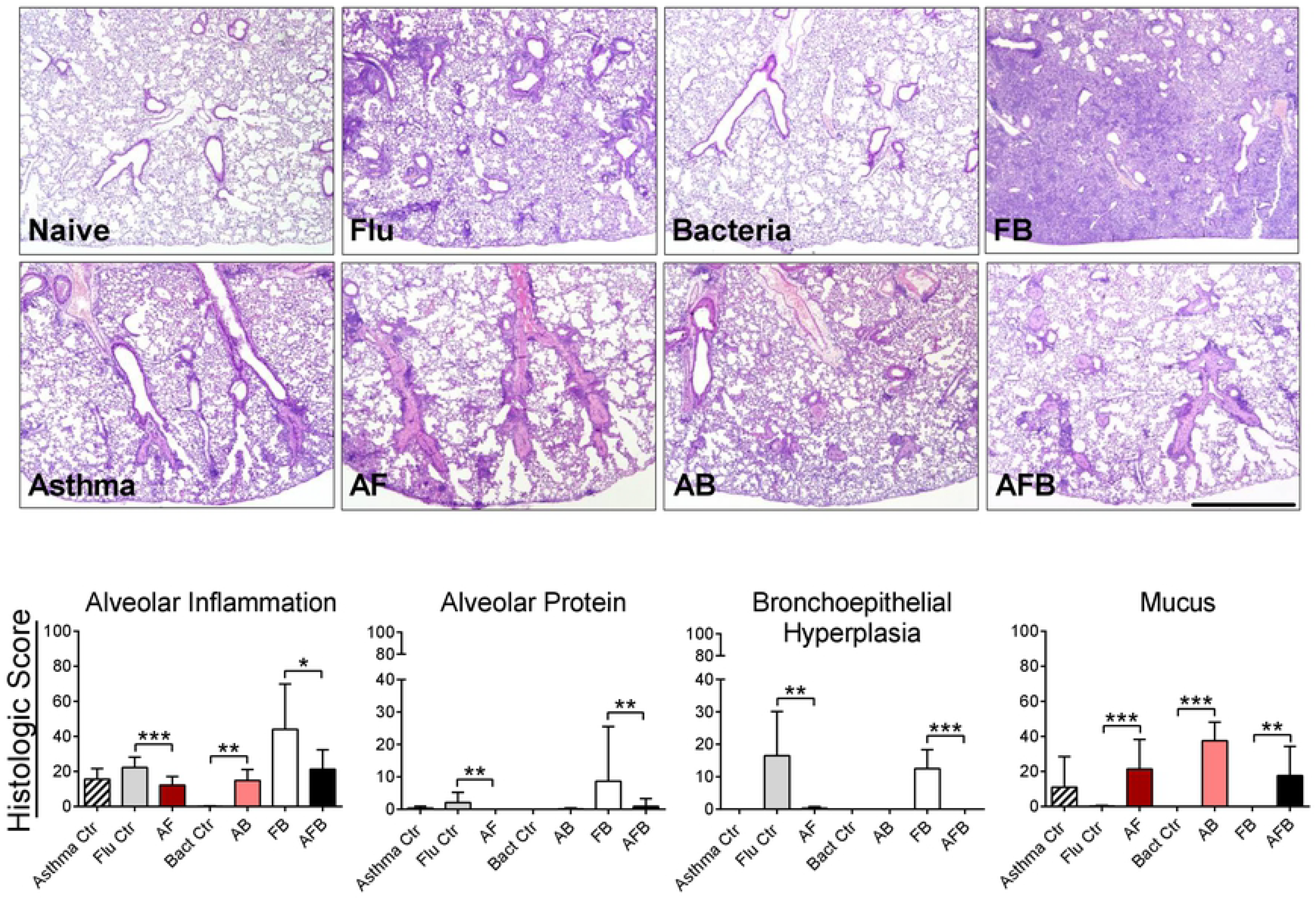
Pulmonary tissue inflammation in triple-disease model. Hematoxylin and eosin stained sections of naïve mice showed no visible inflammation, while mice with asthma showed inflammation around the large airways. Influenza (Flu) virus infected mice had some parenchymal inflammation which was also evident in the asthma and flu (AF) dual disease mice. While no inflammation was evident in the bacteria (Bact) group, allergic mice infected with bacteria (AB) had some inflammatory foci. Co-infection with Flu and Bact (FB) had significant inflammation in the parenchymal spaces in stark contrast to co-infected allergic mice (AFB). While scored criteria of inflammation were reduced in AFB compared to FB, mucus production was maintained at high levels in all allergic mice. Data are representative of one study from two independent studies. n=5-7 mice per group. Data were analysed by Mann-Whitney test against the direct control group for each experimental condition. *P<0.05, **P<0.01, and ***P<0.001.

### Mucosal microbial abundance in the allergic co-infected mice differed from baseline

The importance of the microbiome in health and disease is increasingly recognized (37). Our understanding of the contribution of endogenous microbes in the gastrointestinal system is more advanced than that of the respiratory system, partially due to the comparatively low biomass and sampling difficulties. Therefore, studies that incorporate the microbiome into disease pathogenesis are important. Herein, we investigated the microbial diversity in the BAL and lungs by comparing the microbiome abundance in each niche of each group in comparison to naïve controls at the genus level (Table S1). Differential abundance changes of microbiota under different conditions although the changes were not significant (p>0.05) after multiple test correction across all comparisons.

Comparison of Asthma vs. naïve in BAL, the PCoA ordination based on the abundance profiles did not show a clear separation of sample clusters (**Fig 4A**). *Hyphomicrobiaceae* is enriched in asthma of BAL (p=0.02), while *Methylophilaceae, Clostridiaceae*, and *Mycoplana* were decreased in asthma of BAL (p< 0.05). Clear separation of clusters between Asthma and naïve mouse lungs was evident with enrichment of *Exiguobacterium, Desulfotomaculum* and S24_7 of *Bacteroidales* (p<0.05) and reduction of *Veillonella* (p=0.01) in Asthma (**Fig. 4A**). Slight overlap between Flu-and Bact-controls occurred with naïve microbiome clusters in both niches (**Fig 4B** & **C**). *Rhodocyclaceae, Lysobacter, Skermanella* and *Sphingomonadaceae* were enriched in the BAL of Flu controls while *Mycoplana* and *Cytophagaceae* were depleted (p<0.05). *Lactobacillus, Acetobacteraceae, Balneimonas*, and *Dietzia* were enriched in the lungs of Flu controls (p<0.05) while *Fusobacterium* and *Veillonella* were particularly reduced (p<0.02) (**Fig 4B**). Differentially, *Clostridiales*, S24_7 of *Bacteroidales*, RF39 of *Mollicutes, Capnocytophaga, Roseomonas* of *Acetobacteraceae* were enriched while *Nitrospira* and *Micrococcaceae* were reduced in BAL of mice infected with *Spn* (p<0.05). *Skermanella* dominated in the lungs of Bacteria-only control mice while *Rothia* and *Fusobacteria* were reduced (p<0.01) (**Fig 4C**).

**Figure 4.**
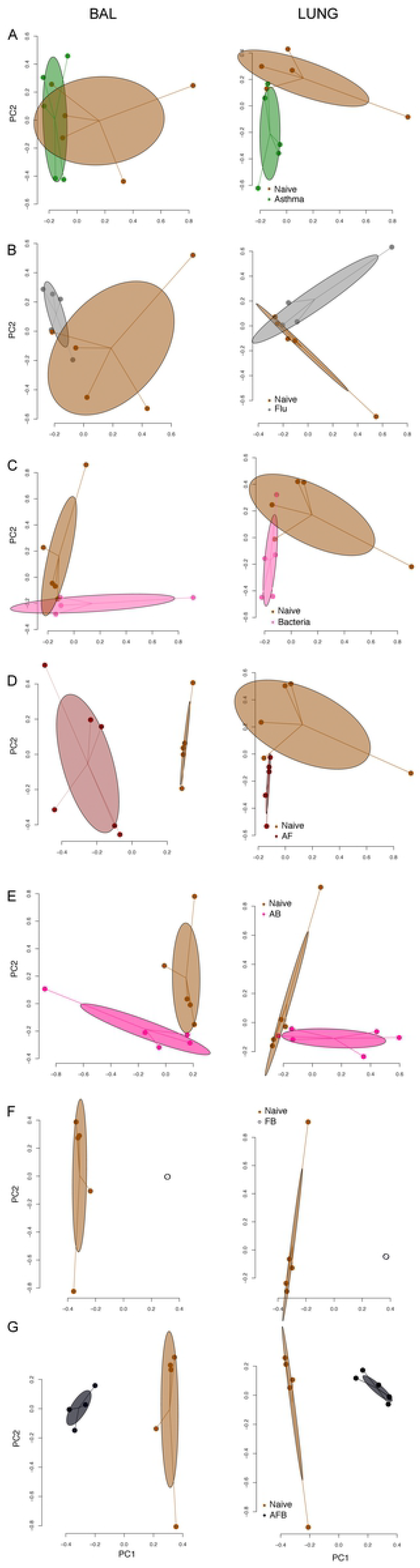
Microbiome profiles of each disease group in comparison with the microbiome profile of naïve mice. Principle coordinate analyses of the microbiome abundance revealed clear separation of clusters in the comparisons of AF in BAL, FB and AFB in both BAL and lung. Data are representative of one study of two independent studies.

Clear separation of clusters were identified between AF and naïve mouse BAL samples wherein *Ruminococcus, Allobaculum, S24_7* of *Bacteroidales* and *Clostridiales* were enriched in AF (p-value < 0.01) and multiple taxa were depleted especially *Ruminococcus* (p=0.005). (**Fig 4D**). Minor overlap was observed between the lung microbiota of AF and naïve mice with *Skermanella, Ruminococcaceae* of *Clostridiales*, S24_7 of *Bacteroidales* and *Nitrospira* highly enriched while *Propionibacterium* and *Corynebacterium* were reduced (p=0.01) in AF group (**Fig. 4D**).

Microbial clusters in the BAL and lungs of AB mice were slightly separated from naïve with enrichment of *Oscillospira* of *Ruminococcaceae* (p<0.01) and depletion of *Nitrospira* (p<0.01) in the AB group (**Fig 4E**). Tight clusters showed clear separation of lung microbiota between the FB group and naïve mice where *Streptococcus* was highly enriched in the BAL (p<0.01) and >20 taxa were depleted in both BAL and lung (p<0.05) in the FB group (**Fig 4F**). Similarly, clear separation of clusters between the AFB group and naïve animals was observed in both niches (**Fig 4G**). *Bacillaceae, Streptococcus, Turicibacter*, and *Bacteroides* were enriched in BAL of AFB (p<0.05) while *Streptococcus* was enriched in AFB lungs (p=0.01) and multiple taxa were reduced in both niches of AFB mice.

### Antibiotic treatment stunted the protection from infection-induced morbidity in allergic mice and worsened influenza morbidity while impacting inflammation

Antibiotic (Abx) overuse is a growing concern with both short-and long-term implications. We hypothesized that a microbiome dysbiosis induced by Abx treatment will increase synergistic pathogenesis of IAV and *Spn* in allergic hosts. Mice were treated daily for two weeks with levofloxacin, a commonly used Abx for respiratory infections, prior to virus infection (**Fig 5A**). Antibiotic-treated IAV-infected mice (Flu ctr) had significantly lower nadir than untreated counterparts (**Fig 5B**). Abx treatment did not alter weight curves in the other groups except the triple-disease state (AFB) in which case allergic co-infected mice treated with antibiotics exhibited weight loss similar to Flu control mice (**Fig 5B**). A concomitant increase in the bacterial burden in the AFB group occurred in all three niches tested while the increase in the FB group did not reach statistical significance in the blood (**Fig 5C**).

**Figure 5.**
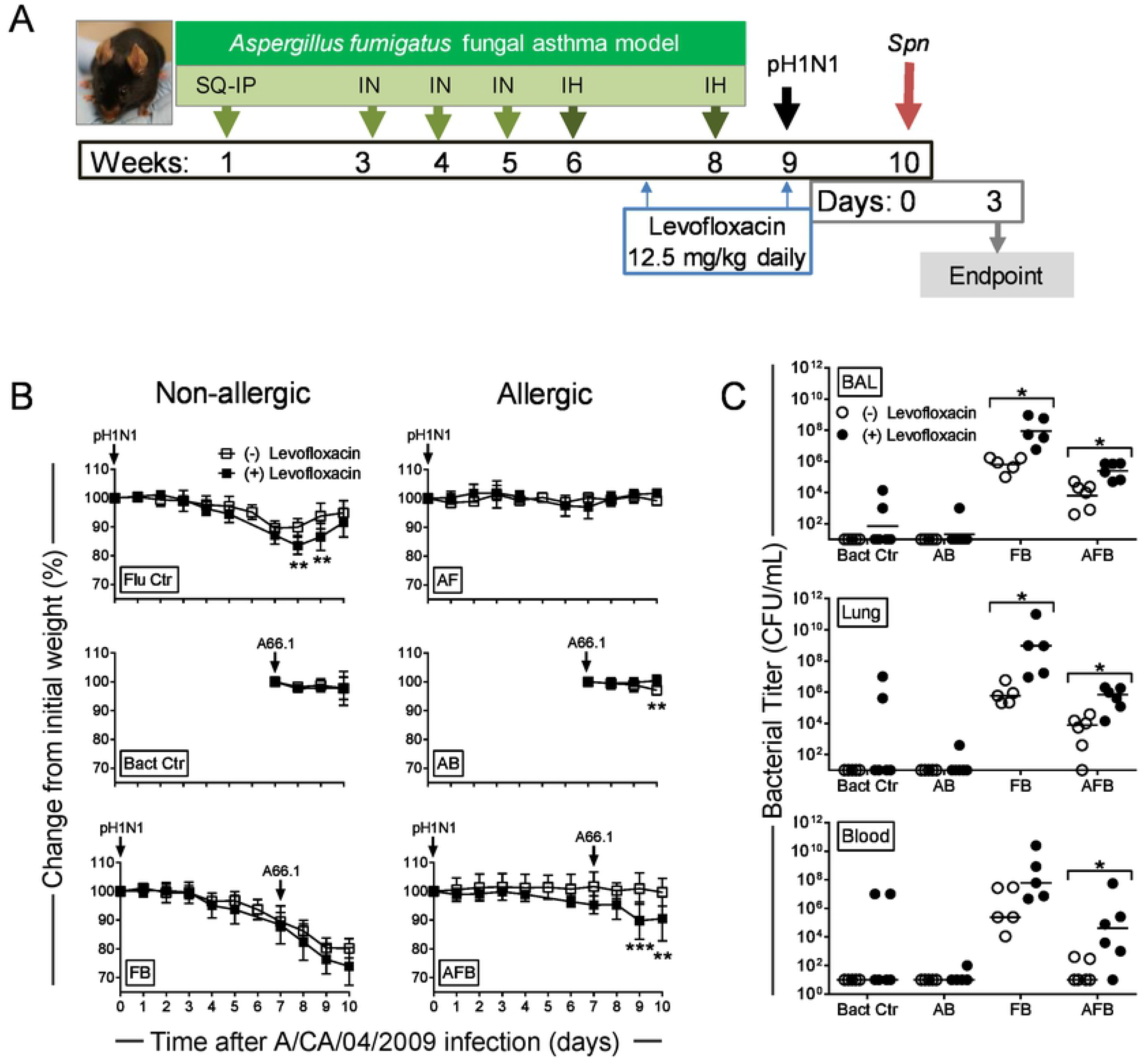
The impact of antibiotic treatment on disease pathogenesis of respiratory infections in allergic hosts. Timeline of model including antibiotic treatments (A). Changes in weight in each group in antibiotic treated mice in comparison to mock-treated groups (B). Bacterial burden in each disease group in comparison to antibiotic treated groups (C). Data are representative of one independent study. n=5-6 mice per group. Data in B were analysed by two-way ANOVA with Sidak’s multiple comparisons test and Data in C were analysed by Mann-Whitney test. *P<0.05, **P<0.01, and ***P<0.001. Ctr: control; F and Flu: influenza virus; B and Bact: bacteria; A: asthma.

While the total cell numbers were reduced in all groups after levofloxacin treatment (**Fig 6A**) compared to untreated mice (as shown in Fig 2), the cell types most dramatically altered were macrophages and CD8^+^T cells (**Fig 6B**). Most cell types were drastically reduced after Abx treatment, while neutrophils were increased in the Flu control and FB groups while CD4^+^T cells and CD8^+^T cells were increased in the Flu control group (**Fig 6B**). Interestingly, eosinophils were reduced in all groups, but most notably in the AB and AFB groups (**Fig 6B**). When we normalized the cell types that were measured in each group, the immune profiles were notably of different compositions between groups (**Fig 6C**) and in comparison untreated mice.

**Figure 6.**
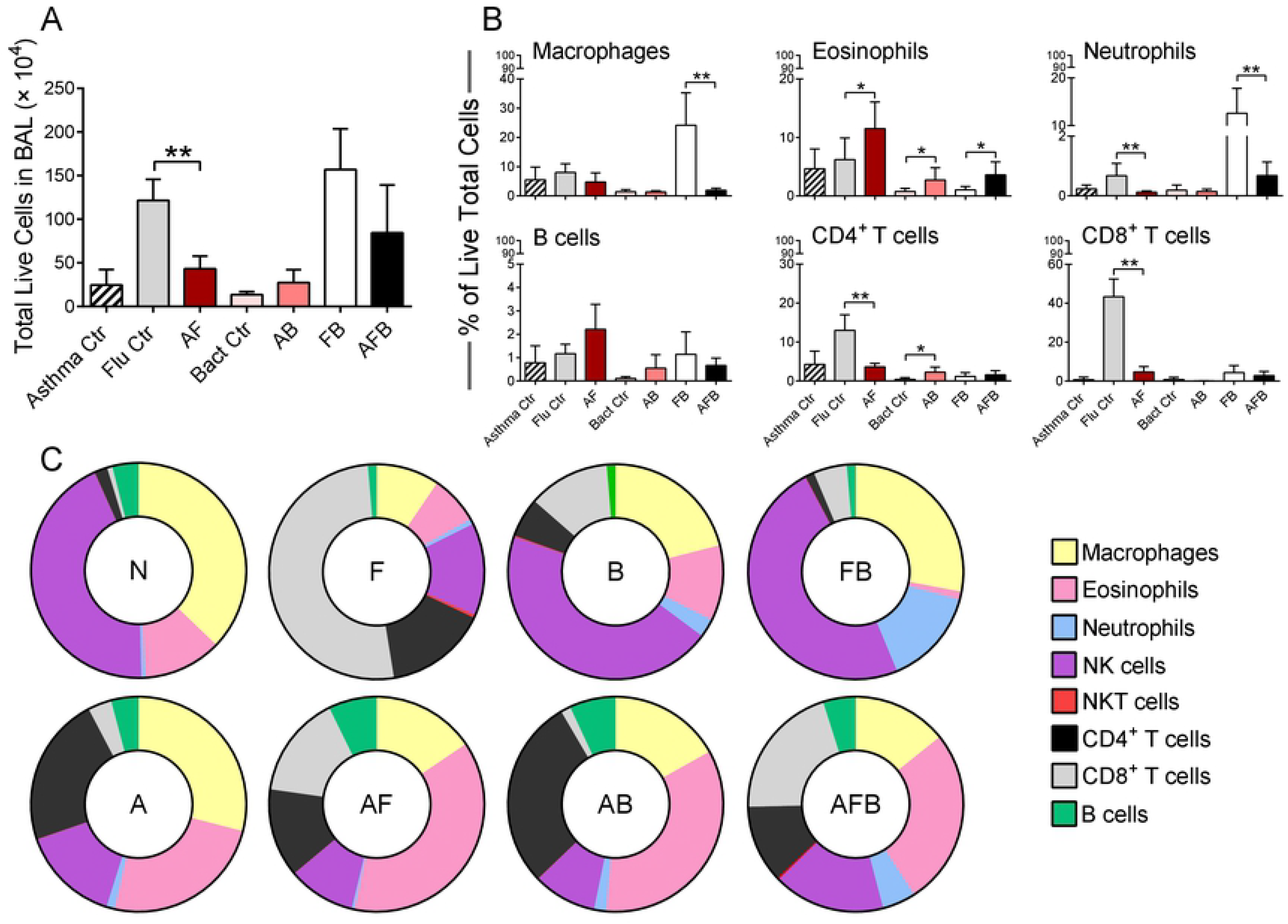
The impact of antibiotic treatment on the immune cell profile in triple-disease model. The number of live leukocytes in the bronchoalveolar lavage (BAL) were enumerated in each group (A) and the various cell populations were identified by flow cytometry (B). Normalized cell populations were used to identify shifts in cell types in each disease state (C). Data are representative of one study from four independent studies. n=5-6 mice per group. Data were analysed by Mann-Whitney test against the direct control group for each experimental condition. *P<0.05 and **P<0.01. Ctr: control; F and Flu: influenza virus; B and Bact: bacteria; A: asthma; N: naïve.

We investigated the pro-inflammatory cytokine milieu in the BAL and the lungs to better understand the immune pressures influencing infiltrating leukocytes and structural cells. In general, the cytokine backdrop in BAL fluid had very little change compared to baseline in all groups except in the FB group (for most analytes) wherein nearly all markers were significantly greater than the AFB group (**Table 1**). Similar trends were observed in the lung homogenates (**Table 1**). Treatment of mice with Abx affected the cytokine profile in both the BAL and lungs with significant increases occurring in both the FB and AFB groups (**Table 1**). However, even after Abx treatment, most cytokines were significantly more highly expressed in the FB compared to the AFB group (**Table 1**).

**Table 1.**
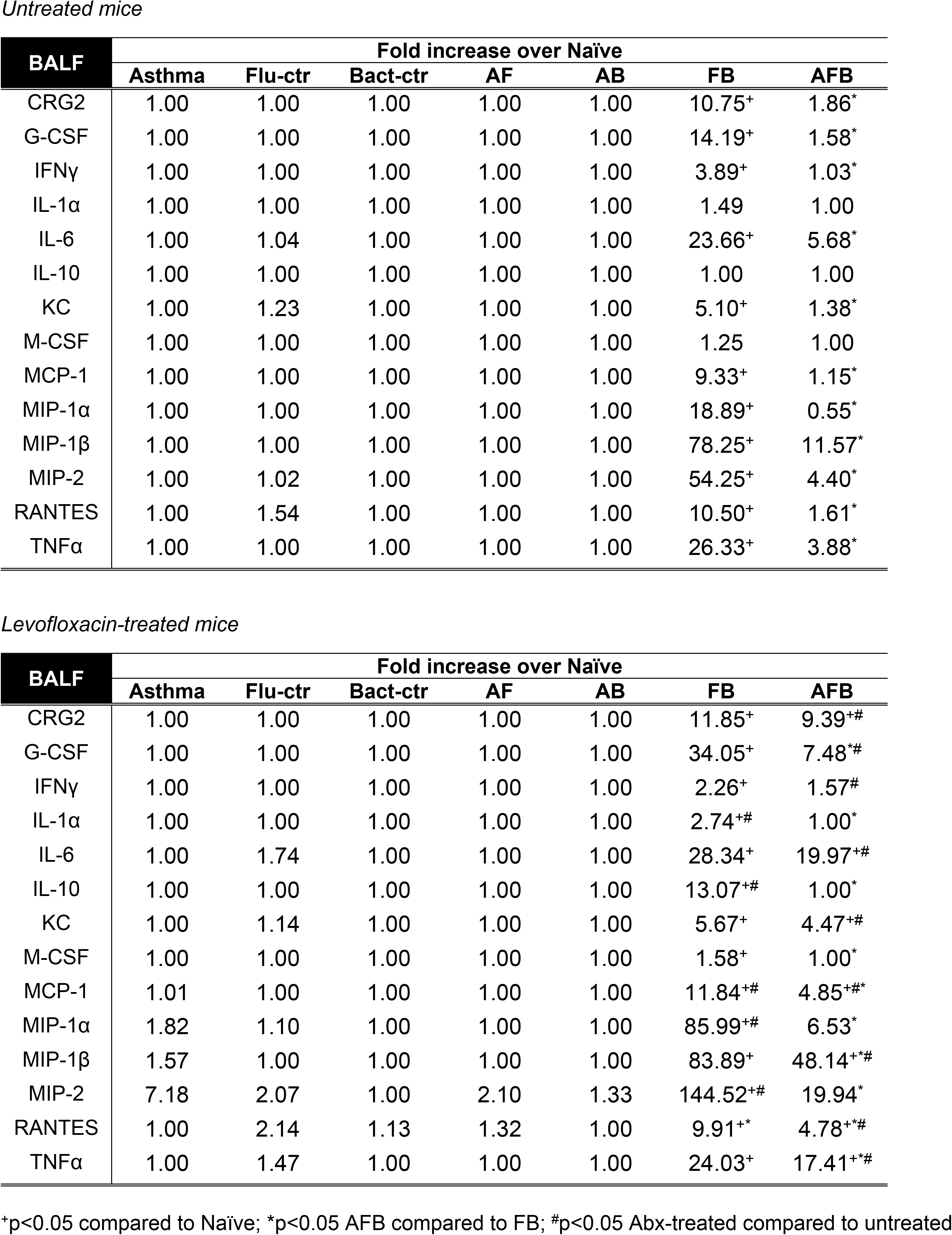
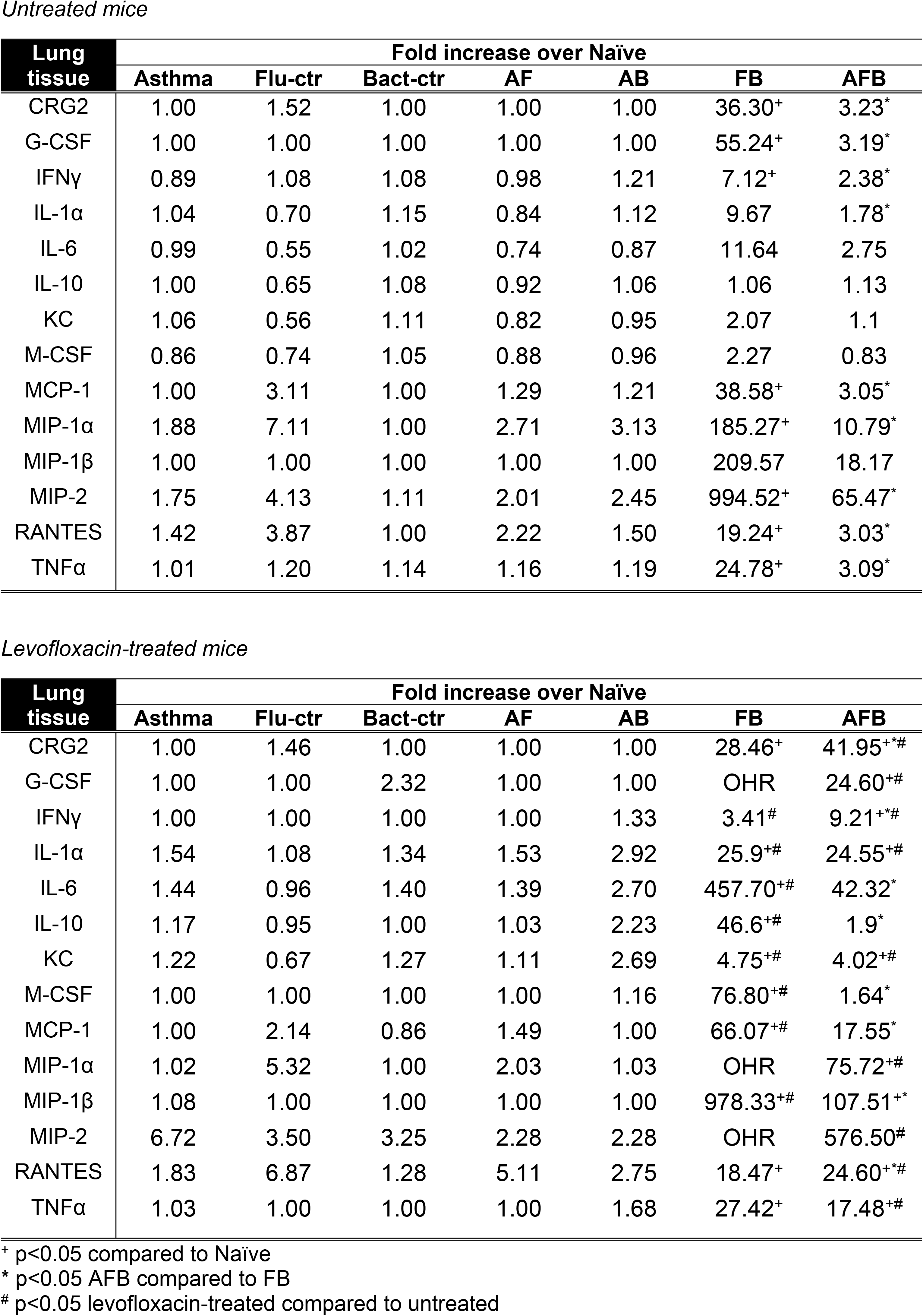
Mucosal Cytokines in Mouse Bronchoalveolar Lavage Fluid (BALF) and Lung Homogenate.

### Mucosal microbiome diversity was reduced by antibiotic treatment

In order to determine if we were successful in inducing dysbiosis, we analysed the microbiome in the BAL and lungs of mice that were treated with Abx in comparison to untreated mice. The PCoA ordination indicated that microbiota of Abx-treated mice formed independent clusters from untreated mice except in the AFB group (**Fig 7**). Differential abundance analyses indicated that the majority of treated mice had reduced taxa abundance. While *Facklamia, Bacillaceae*, and *Enterococcus* (p=0.046) were enriched in BAL of Abx-treated Asthma group, significant depletions in were identified in multiple genera especially *Alphaproteobacteria* and *Actinobacteria* (Table S2). Distinct clusters in the lungs of asthma group (**Fig 7A**) showed significant enrichment for *Anaerococcus* of *Clostridia* and *Lactobacillus* in the Abx-treated group, while numerous genera including *Proteobacteria, Firmicutes* and *Actinobacteria*, had decreased abundance after Abx treatment.

**Figure 7.**
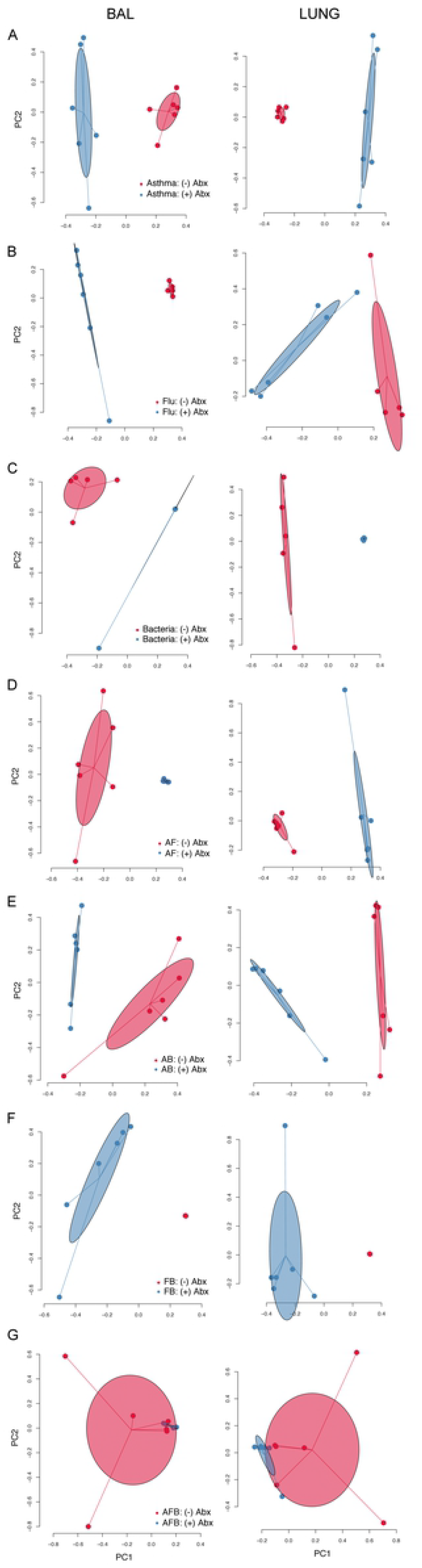
Microbiome profiles of each disease group with and without antibiotics (Abx). Principle component analysis of Abx-treated mice in each group (blue) compared to untreated mice (red) showed independent clustering of microbiome for all study groups except in the AFB. Data are representative of one study.

A majority of taxa in the microbiome of Abx-treated Flu control mice were reduced except for a few genera (*Peptoniphilus, Selenomonas*, and *Enterococcus*) that were enriched in the BAL clusters (**Fig 7B**). *Fusobacterium* enriched the lung microbiome cluster of Abx-treated Flu controls while *Streptococcus* enriched both BAL and lung. All identified taxa in the microbiota of Bacteria-only controls had a significantly reduced abundance after Abx-treatment except for *Streptococcus* that was enriched in the lungs with the clearly separated clusters (**Fig 7C**).

Individual clusters were observed between Abx-treated and untreated AF group (**Fig 7D**), with *Streptococcus* (p=0.03) and *Bradyrhizobium* (p=0.04) enrichment in BAL and *Leptotrichia* in the lung (p=0.04) of treated AF mice. Multiple genera including *Proteobacteria, Firmicutes* and *Actinobacteria* were reduced in the BAL of Abx-treated AB mice while *Lactococcus* and *Sphingopyxis* were enriched. Similarly separated clusters in the AB group lungs (**Fig 7E**) had dynamic enrichment/depletion in *Firmicutes, Fusobacteria* and *Proteobacteria* after Abx-treatment.

Clearly separated clusters were evident between the Abx-treated and untreated FB group in both niches (**Fig 7F**). Interestingly, the majority of taxa have increased abundance with *Streptococcus* being the only taxa with decreased abundance after treatment in both BAL and lung in FB group. Overlapping clusters between Abx-treated and untreated mouse microbiota in the BAL and lungs were only observed in the AFB group (**Fig 7G**) wherein most of the identified taxa had reduced abundance; the only exceptions were *Streptococcus* and *Anaerococcus* that were enriched after antibiotic treatment in both niches.

## Discussion

Susceptibility to and severity of respiratory infections are dictated by a multitude of variables from the perspective of both host and pathogen. Numerous predisposing conditions have been described in having both protective and detrimental roles in the susceptibility of patients to respiratory infections. Such underlying conditions include asthma (27, 38), obesity (39, 40), malnutrition (41), impaired liver function (42), and sickle cell disease(43-45). From the standpoint of respiratory pathogens, the disease strategies for two of the major etiological agents, IAV and *Spn*, are well described in terms of the main virulence factors required during infection of the respiratory tract (46-50). Since virulence strategies utilized by pathogens vary considerably based on the host environment, underlying conditions present unique challenges and opportunities for the invading pathogens. The triple-disease murine model utilized here is clinically relevant as asthmatics are considered at risk for respiratory infections and functions as tool with which to investigate complex host-pathogen interactions within the allergic host.

Predisposing host conditions provide a unique environment in terms of nutrient bioavailability, receptor expression, and inflammatory status that can alter the virulence strategies of potential pathogens. Influenza synergy with secondary pneumococcal pneumonia is one of the best characterized exacerbations of respiratory infection, with both clinical data and murine models supporting this viral-mediated superinfection (36). Somewhat counterintuitively, our data show that the well-established detrimental synergy between IAV and *Spn (11, 13, 26, 36, 51)* is reduced in the context of experimental allergic asthma where IAV+ *Spn* co-infection in allergic mice resulted in a dramatically altered inflammatory landscape correlating with less overall lung inflammation and damage, reduced morbidity and enhanced bacterial clearance. It has been previously suggested that a rapidly induced short lasting inflammation may hinder IAV pathogenesis, reduce host pathology (52, 53), and beneficial during IAV-*Spn* co-infection (54). Our data suggest that allergic hosts may hold a similar advantage when infections occur during heightened allergic inflammation.

It is becoming increasingly evident that resident bacterial species within the respiratory tract can alter infection susceptibility to exogenous viral and bacterial pathogens (55-58). Perturbations to the respiratory microbiome may occur during disease states as well as during Abx treatments (59, 60). Early life exposures to microbial agents have been demonstrated both as triggers (61) and inhibitors (62) of asthma development based on the agent, atopy, and age (63-65). Early Abx use associated with allergic asthma in young children even when accounting for bias inherent from when Abx are commonly prescribed to treat early symptoms of asthma (66, 67). Analysis of microbial communities in the respiratory tract revealed the asthmatic host to have a distinct microbial signature compared to healthy controls, suggesting these alterations in the respiratory microbiome may contribute to the protective capacity of asthma during IAV+*Spn* co-infection. If this mechanism was operative, we hypothesized that perturbation of the microbial communities would diminish the protective capacity of allergic asthma in the context of influenza-pneumococcal co-infection. This was accomplished by Abx administration, which significantly altered the microbial composition of the respiratory tract. Concurrent with this perturbation of the microbial communities was a significant decrease in the protective capacity allergic asthma engenders against IAV+*Spn* co-infection. Similar reductions in host protection occur after perturbations to gut microbiome with Abx (68, 69). However, since microbiome disruption with Abx also resulted in a modification to the immune cell profile (with a notable reduction in macrophages and increase in pro-inflammatory cytokines), increased susceptibility to co-infection in the allergic mice may have resulted from a decrease in immunity. While this ‘chicken or egg’ paradox is currently under investigation by our group, it is clear that both the microbiome and immune system play a crucial role in mediating host protection during complicated respiratory infections, and that Abx should be prescribed with caution, especially in patients with chronic underlying conditions as microbiome and immune baselines are likely drifted from an otherwise healthy host.

The extensive alterations in the respiratory tract during allergic asthma encompass both immunological and microbiological differences that can have profound impact on susceptibility to infection. These data suggest that allergic asthma may provide a significant advantage in terms of morbidly and mortality against certain types of respiratory infections and that this protection is in partly mediated by alterations in the resident flora and immune milieu. Due to the complex interplay between the microbiota and host immunity, there are likely additional host signaling pathways important for the modulation of the sensitivity of the allergic host to infection. The observation that Abx partially ameliorated this protection also underscores the importance of appropriate Abx use, as prior Abx exposure enhances the susceptibility of mice with allergic asthma to subsequent infection. This is of particular importance to asthmatic patients, as the early symptoms of asthma are similar to respiratory infection, potentially resulting in inappropriate Abx usage (70). These data underscore the complex and interwoven relationship between host immune status and the resident bacterial flora in terms of infection susceptibility.

## Methods

### Pathogen strains and growth

A clinical Influenza A Virus isolate recovered during the 2009 influenza A virus pandemic (A/CA/04/2009) was gifted by Dr. Webby (St. Jude Children’s Research Hospital, Memphis, TN), propagated in Madin-Darby canine kidney (MDCK) cells (ATCC, Manassas, VA), sequence verified to be void of mutations in HA and NA genes, and stored as single use aliquots at −80°C. The concentration of virus was determined by using the TCID50 method in MDCK cells and virus was diluted in sterile PBS to desired concentration for mouse inoculations.

*S. pneumoniae* A66.1L is a type 3 strain constitutively expressing luciferase (71), gifted to us by Dr. Jon McCullers (University of Tennessee Health Science Center, Memphis, TN). A66.1L was cultured in Todd-Hewitt broth (Difco Laboratories, Detroit, MI) supplemented with yeast extract (ACROS Organics, NJ) to OD620 0.15 corresponding to log growth phase. Cultures were centrifuged at 2671 *g*/4°C, re-suspended in fresh Todd-Hewitt Yeast broth supplemented with 30% autoclaved glycerol, then frozen in single use aliquots at −80°C. Aliquot concentration was determined by enumerating colony growth on agar plates with 5% sheep blood (Remel, Thermo Fisher, Lenexa, KS) at the time of freezing, and confirmed by plating at least one week after storage at −80°C. To prepare mouse inocula, A66.1L was thawed at room temperature and serially diluted in PBS in 15 mL tubes, then vortexed. Inocula concentration was confirmed by colony growth on blood agar plates.

### Ethics Statement

Animal research described was approved by the University of Tennessee Health Science Center’s Institutional Animal Use and Care Committee (IACUC) under protocol 15-003.0 and 18-003.0 to be in accordance with the Office of Laboratory Animal Welfare guidelines.

### Animals and housing conditions

No discernible differences in gender were noted after subjecting mice to the fungal asthma and influenza model, female mice were used in these studies. Six week-old female C57BL/6J mice were purchased from Jackson Laboratories (Bar Harbor, ME) and maintained in sterile microisolator cages on α-dri bedding within the animal biosafety level-2 facilities at SJCRH and University of Tennessee Health Science Center for 1 week prior to being used in experiments. The animal housing facilities were on a 12 h light-dark cycle and all work with animals was done during the light cycle. Animals were fed autoclaved chow and provided autoclaved water in bottles *ad libitum*.

### Mouse fungal asthma model

Mice were subjected a previously described and characterized model of *Aspergillus fumigatus*-induced allergic asthma (33, 72). In brief, whole *A. fumigatus* extract (Greer Labs, Lenoir, NC) was used to sensitize mice over a period of five weeks prior to subjecting them to airborne dry conidia in a nose-only inhalation chamber for 10-minutes two weeks apart (25). Allergen sensitized and challenged mice were referred to as the “Asthma Ctr” group.

### Animal infections

Mice were lightly anesthetized with isoflurane and intranasally infected with 1000 TCID50 A/CA/04/2009, diluted in 50 μL PBS. Seven days after the viral infection, mice were anesthetized with isoflurane and intranasally infected with 600 colony forming units (CFU) of A66.1L in 100 μL PBS. Each mouse was weighed prior to challenge and every 24 hours after until harvest to monitor weight change. Animals that lost more than 30% of their starting body weight, or exhibited severe signs of morbidity, were euthanized for ethical reasons and recorded as having died on that day. Each group of mice that were infected with a single agent were named after the pathogen as influenza virus (Flu Ctr) and bacteria (Bact Ctr) only. Naïve (N) mice that had no treatments served to determine baseline immune and microbiome information. Allergic mice that were infected with influenza (Flu) virus (23) were referred to as the “Asthma and Influenza” (AF) group and those additionally infected with bacteria were considered the “Asthma, Influenza, and Bacteria” (AFB) group. Allergic mice that were infected with bacteria were referred to as the “Asthma and Bacteria” (AB) group.

### Antibiotic treatments

Naïve and mice in the various treatment groups were intraperitoneally administered 12.5 mg/kg levofloxacin (Akorn Inc., Lake Forest, IL) daily at the equivalence of one week after the first allergen challenge and ending on the day of viral infection. No visible differences occurred during disease pathogenesis in mice after antibiotic treatment.

### Bioluminescence measurements

The use of the A66.1L strain allowed for the visualization of bacteria during the active infection in mice. We measured bioluminescence by a Lumina IVIS CCD camera (Perkin Elmer, Waltham, MA). Images were processed with Living Image software, version 4.5.5. Settings for image acquisition were a one second photograph following by one minute luminescence measurement with the bining set at four.

### Tissue harvest and bacterial enumeration

Tissues were harvested within a class 2A biosafety cabinet with strict adherence to aseptic technique to ensure that samples were not contaminated by exposure to environmental agents. Bronchoalveolar lavage (BAL) was performed using two consecutive infusions of 1 mL sterile PBS. BAL cells were cytospun and stained with Diff-quik (StatLab, McKinney, TX) for visualization. Cells were, then centrifuged at 4°C and BAL fluid (BALF) was stored at −80°C until use. Red blood cells in the cells pellet were removed by lysis with cold water and resultant cells enumerated on a Countess Automated Cell Counter (Invitrogen, Carlsbad, CA) then stained for flow cytometry.

Blood was recovered from the chest cavity, and serum was separated by centrifugation and stored at −80°C. Lungs were excised and homogenized in 1 mL PBS containing cOmplete Proteinase Inhibitor Cocktail (Roche Diagnostics, Mannheim, Germany) then centrifuged at 4°C to remove cell debris. Cell-free supernatant was stored at −80°C until cytokine quantitation. In some experiments, spleen, mediastinal lymph nodes and left lung lobes were perfused with 10% neutral-buffered formalin and processed for hematoxylin and eosin staining.

Whole lung homogenate, whole blood, and BAL samples were serially diluted in sterile PBS and 10 μL of dilutions were plated on blood agar plates. Plates were incubated at 37°C with 5% CO2 for 12-13 hours, and colony growth was enumerated.

### Histopathologic analysis

Lungs were first infused and then immersion-fixed in 10% neutral buffered formalin before processing and embedding in paraffin. Tissue sections were stained with hematoxylin and eosin, and the severity and extent of specific pulmonary lesions such as interstitial and alveolar inflammation, alveolar protein exudate, hyaline membrane formation, septal thickening, epithelial hypeplasia, and denuded bronchioles, were assessed and graded in a blinded manner by a veterinary pathologist. Separate severity grades for each type of lesion were assigned as follows: 0, no lesions detected; 1, minimal, rare, barely detectable lesions; 2, mild multifocal, small focal, or widely separated lesions; 3, moderate, multifocal, and prominent lesions; 4, marked, extensive-to-coalescing areas; and 5, severe and extensive lesions with pulmonary consolidation. These severity grades were then converted to weighted semi-quantitative scores as follows: 0 = 0; 1 = 1; 2 = 15; 3 = 40; 4 = 80; and 5 = 100.

### Flow cytometric analysis

Cells in the airways were identified using flow cytometry. BAL cells were incubated with human γglobulin to prevent nonspecific binding of antibodies to Fc-receptor, then stained with fluorescently tagged antibodies. Samples were fixed with BD Biosciences stabilizing fixative and stored in the dark at 4°C until acquisition on a LSR Fortessa (BD Biosciences, San Jose, CA). Data were analyzed using FlowJo v 10.1r5 (FlowJo LLC, Ashland, OR). Unstained cells, single-color controls and isotype controls were used for instrument settings and compensation. Abs were purchased from BD Biosciences unless specified otherwise. Antibodies used in this study include: CD19-PerCP/Cy5.5 (1D3), CCR3-A647 (83103), CD3e-PE/Cy7 (145-2C11), CD4-A700 (RM4-5), CD8a-FITC (53-6.7), Ly6G-V450 (1A8), Mac3-Biotin (ebioABL-93, eBioscience|Thermo Fisher, Waltham, MA), NK1.1-APC/Cy7 (PK136), NP-dextramer-PE (Immudex, Fairfax, VA), Siglec-F-PE-CF594 (E50-2440), Streptavidin-BV605 (BioLegend) and matching isotypes control antibodies.

### Cytokine quantitation

Cytokines present in the airways and interstitial lung tissue were quantitated using magnetic Luminex multiplex assays (R&D) according to the manufacturer’s instructions. BALF and cell-free lung homogenates were stored at −80°C in single-use aliquots after harvest until cytokine assay, and used undiluted or diluted ½ in Calibrator Diluent (R&D Systems (Minneapolis, MN), respectively. Beads were read using a Luminex MagPix (R&D Systems). Analyte concentration was determined using MagPix software xPONENT 4.2. Samples which had a reading below the limit of detection were assigned the value of the limit of detection. The mean concentration for the group for each analyte was used to calculate the fold change compared to the mean concentration of the same analyte in the naïve group.

### 16S sequence analysis

BAL and whole lung homogenates were centrifuged at 500 ×*g* at 4°C to remove cell debris, then subsequently centrifuged at 2671 ×*g* at 4°C to pellet bacteria in the samples. Bacterial pellets were re-suspended in 50 μL of BALF or lung homogenate supernatant and stored at −80°C until DNA extraction. Samples were thawed on ice and DNA was extracted using FastDNA-96 Soil Microbe DNA kit (MP Biomedicals, Santa Ana, CA) according to the manufacturer’s instructions, and a FastPrep-96 instrument (MP Biomedicals). Extracted nucleic acid was stored at −80°C until use.

The V1-V3 variable region of the bacterial 16S gene was amplified using the Bioo NEXTflex 16S amplicon library preparation kit. The multiplexed products were then subsequently utilized for high-depth sequencing on the Illumina Mi-seq platform to obtain comprehensive relative abundance of the bacterial composition at the respective time points with 300 bp paired end read lengths. The quality of the raw 16S rRNA pair-ended reads are initially examined by FastQC.(73) Low quality reads and bases are trimmed by Trim Galore (74). The reads are then merged by PANDAseq(75) and subsequently processed by QIIME (76). The closed reference mapping protocol of QIIME was used for OTU assignment. Specifically, the read sequences were clustered into Operational Taxonomic Units (OTUs) at 97% sequence similarity using the UCLUST algorithm (77). A representative sequence was then selected from each OTU for taxonomic assignment using the Greengenes database (78) as the reference. The alpha diversity estimates were calculated using the R Phyloseq package (79). Kruskal-Wallis non-parametric tests was used to test the significance of the diversity difference. The linear discriminant analysis (LDA) effect size (LEfSe) method was used to test the significant difference of relative abundance of taxa among groups (80).

## Acknowledgments

The authors would like to thank the Animal Resource Centers at St. Jude Children’s Research Hospital and University of Tennessee Health Science Center for husbandry and care of animals during the course of these studies. We would also like to thank Yanyan Lin, Hannah Buser, and Laura Doorley all formerly at the Children’s Foundation Research Institute for help in tissue harvests and processing. The studies described herein were partially funded through the Young Investigator Award (Le Bonheur Children’s Foundation) and the NIH, R01-AI125481 to AES. JWR is supported by NIH, U01-AI124302 and R01-AI110618 and by the American Lebanese Syrian Associated Charities (ALSAC).

## Author Contributions

KSL performed infections, harvested tissues, determined bacterial titers, performed flow cytometry and analysis, extracted DNA for 16S sequencing and multiplex for cytokines. ARI performed pilot studies to determine optimum bacterial infections, harvested tissues, determined bacterial burden by IVIS CCD camera, and performed qPCR for BAL and lung samples prior to genomic sequencing for microbiome analyses. T-CC performed data analyses for microbiome. MP subjected mice to the fungal asthma model, helped with tissue harvest, treated mice with antibiotics, and performed viral titrations. PV performed histopathologic analyses. JWR designed the microbiome and antibiotic treatment studies. AES designed and directed the studies, analysed data, and created Figures. KSL, JWR, and AES co-wrote the first draft of the manuscript, and all authors edited and approved the final version.

## Competing Interests

The authors have no conflicts of interest to declare.

## Supporting Information Legends

**Supplemental Table 1.** Bacterial classifications based on 16S sequence analysis for lungs and BAL from A, AF, AB, AFB, F, and B groups.

**Supplemental Table 2.** Bacterial classifications based on 16S sequence analysis for lungs and BAL from A, AF, AB, AFB, F, and B groups from mice on levofloxacin prophylaxis.

**Table**

**Table 1. Fold change in mucosal cytokines in mice subjected to allergen and infectious agents.**

Cytokine levels in the bronchoalveolar lavage fluid (BALF) and lungs were determined for each group and used to determine the change over baseline (naïve) levels in mice that were treated with antibiotics and those that were not treated. Data were analysed by two-way ANOVA with Dunnett’s multiple comparisons test. F and Flu: influenza; B and Bact: bacteria; A: asthma.

## References

1. Association AL. Estimated Prevalence and Incidence of Lung Disease by Lung Association Territory. 2014.

2. Association AL. Asthma and Children Fact Sheet [Website]. 2012 [cited 2014 06/24/2014]. Available from: http://www.lung.org/lung-disease/asthma/resources/facts-and-figures/asthma-children-fact-sheet.html.

3. CDC. Asthma in the US [Website]. 2011 [updated 05/03/2011; cited 2014 06/23/2014]. Statistics on asthma in the US]. Available from: http://www.cdc.gov/vitalsigns/asthma/.

4. Pelaia G, Vatrella A, Gallelli L, Renda T, Cazzola M, Maselli R, et al. Respiratory infections and asthma. Respiratory medicine. 2006 May;100(5):775-84. PubMed PMID: 16289785.

5. Jackson DJ, Sykes A, Mallia P, Johnston SL. Asthma exacerbations: origin, effect, and prevention. The Journal of allergy and clinical immunology. 2011 Dec;128(6):1165-74. PubMed PMID: 22133317.

6. Juhn YJ, Kita H, Yawn BP, Boyce TG, Yoo KH, McGree ME, et al. Increased risk of serious pneumococcal disease in patients with asthma. The Journal of allergy and clinical immunology. 2008 Oct;122(4):719-23. PubMed PMID: 18790525. Pubmed Central PMCID: 2811957.

7. Talbot TR, Hartert TV, Mitchel E, Halasa NB, Arbogast PG, Poehling KA, et al. Asthma as a risk factor for invasive pneumococcal disease. The New England journal of medicine. 2005 May 19;352(20):2082-90. PubMed PMID: 15901861.

8. Ferkol T, Schraufnagel D. The global burden of respiratory disease. Annals of the American Thoracic Society. 2014 Mar;11(3):404-6. PubMed PMID: 24673696.

9. CDC. Pneumococcal Disease 2015 [cited 2017 Apr 27]. Available from: https://www.cdc.gov/pneumococcal/clinicians/clinical-features.html.

10. CDC. Influenza (Flu). 2017.

11. McCullers JA. Insights into the interaction between influenza virus and pneumococcus. Clinical microbiology reviews. 2006 Jul;19(3):571-82. PubMed PMID: 16847087. Pubmed Central PMCID: 1539103.

12. File TM, Jr. Streptococcus pneumoniae and community-acquired pneumonia: a cause for concern. The American journal of medicine. 2004 Aug 2;117 Suppl 3A:39S-50S. PubMed PMID: 15360096.

13. Taubenberger JK, Morens DM. 1918 Influenza: the mother of all pandemics. Emerging infectious diseases. 2006 Jan;12(1):15-22. PubMed PMID: 16494711. Pubmed Central PMCID: 3291398.

14. Gill JR, Sheng ZM, Ely SF, Guinee DG, Beasley MB, Suh J, et al. Pulmonary pathologic findings of fatal 2009 pandemic influenza A/H1N1 viral infections. Archives of pathology & laboratory medicine. 2010 Feb;134(2):235-43. PubMed PMID: 20121613. Pubmed Central PMCID: 2819217.

15. Weinberger DM, Simonsen L, Jordan R, Steiner C, Miller M, Viboud C. Impact of the 2009 influenza pandemic on pneumococcal pneumonia hospitalizations in the United States. The Journal of infectious diseases. 2012 Feb 1;205(3):458-65. PubMed PMID: 22158564. Pubmed Central PMCID: 3276240.

16. Jain S, Kamimoto L, Bramley AM, Schmitz AM, Benoit SR, Louie J, et al. Hospitalized patients with 2009 H1N1 influenza in the United States, April-June 2009. The New England journal of medicine. 2009 Nov 12;361(20):1935-44. PubMed PMID: 19815859.

17. Gilca R, De Serres G, Boulianne N, Ouhoummane N, Papenburg J, Douville-Fradet M, et al. Risk factors for hospitalization and severe outcomes of 2009 pandemic H1N1 influenza in Quebec, Canada. Influenza Other Respi Viruses. 2011 Jul;5(4):247-55. PubMed PMID: 21651735.

18. Bramley AM, Dasgupta S, Skarbinski J, Kamimoto L, Fry AM, Finelli L, et al. Intensive care unit patients with 2009 pandemic influenza A (H1N1pdm09) virus infection-United States, 2009. Influenza and other respiratory viruses. 2012 Nov;6(6):e134-42. PubMed PMID: 22672249. Pubmed Central PMCID: 4941711.

19. Louie JK, Acosta M, Winter K, Jean C, Gavali S, Schechter R, et al. Factors associated with death or hospitalization due to pandemic 2009 influenza A(H1N1) infection in California. Jama. 2009 Nov 4;302(17):1896-902. PubMed PMID: 19887665.

20. Van Kerkhove MD, Vandemaele KA, Shinde V, Jaramillo-Gutierrez G, Koukounari A, Donnelly CA, et al. Risk factors for severe outcomes following 2009 influenza A (H1N1) infection: a global pooled analysis. PLoS medicine. 2011 Jul;8(7):e1001053. PubMed PMID: 21750667. Pubmed Central PMCID: 3130021.

21. Veerapandian R, Snyder JD, Samarasinghe AE. Influenza in Asthmatics: For Better or for Worse? Frontiers in immunology. 2018;9:1843.

22. Myles P, Nguyen-Van-Tam JS, Semple MG, Brett SJ, Bannister B, Read RC, et al. Differences between asthmatics and nonasthmatics hospitalised with influenza A infection. The European respiratory journal. 2013 Apr;41(4):824-31. PubMed PMID: 22903963. Pubmed Central PMCID: 3612580.

23. Samarasinghe AE, Woolard SN, Boyd KL, Hoselton SA, Schuh JM, McCullers JA. The immune profile associated with acute allergic asthma accelerates clearance of influenza virus. Immunology and cell biology. 2014 Jan 28;92:449-59. PubMed PMID: 24469764.

24. McKenna JJ, Bramley AM, Skarbinski J, Fry AM, Finelli L, Jain S, et al. Asthma in patients hospitalized with pandemic influenza A(H1N1)pdm09 virus infection-United States, 2009. BMC infectious diseases. 2013;13:57. PubMed PMID: 23369034. Pubmed Central PMCID: 3585510.

25. Samarasinghe AE, Hoselton SA, Schuh JM. The absence of the VPAC(2) receptor does not protect mice from Aspergillus induced allergic asthma. Peptides. 2010 Jun;31(6):1068-75. PubMed PMID: 20226823. Pubmed Central PMCID: 2873113.

26. McCullers JA, Rehg JE. Lethal synergism between influenza virus and Streptococcus pneumoniae: characterization of a mouse model and the role of platelet-activating factor receptor. The Journal of infectious diseases. 2002 Aug 01;186(3):341-50. PubMed PMID: 12134230.

27. Juhn YJ. Risks for infection in patients with asthma (or other atopic conditions): is asthma more than a chronic airway disease? The Journal of allergy and clinical immunology. 2014 Aug;134(2):247-57; quiz 58-9. PubMed PMID: 25087224. Pubmed Central PMCID: 4122981.

28. Samarasinghe AE, Melo RC, Duan S, LeMessurier KS, Liedmann S, Surman SL, et al. Eosinophils Promote Antiviral Immunity in Mice Infected with Influenza A Virus. Journal of immunology. 2017 Apr 15;198(8):3214-26. PubMed PMID: 28283567. Pubmed Central PMCID: 5384374.

29. Morens DM, Taubenberger JK, Fauci AS. Predominant role of bacterial pneumonia as a cause of death in pandemic influenza: implications for pandemic influenza preparedness. The Journal of infectious diseases. 2008 Oct 1;198(7):962-70. PubMed PMID: 18710327. Pubmed Central PMCID: 2599911.

30. Saturni S, Contoli M, Spanevello A, Papi A. Models of Respiratory Infections: Virus-Induced Asthma Exacerbations and Beyond. Allergy, asthma & immunology research. 2015 Nov;7(6):525-33. PubMed PMID: 26333698. Pubmed Central PMCID: 4605924.

31. Busse WW, Lemanske RF, Jr., Gern JE. Role of viral respiratory infections in asthma and asthma exacerbations. Lancet. 2010 Sep 4;376(9743):826-34. PubMed PMID: 20816549. Pubmed Central PMCID: 2972660.

32. Kraft M. The role of bacterial infections in asthma. Clinics in chest medicine. 2000 Jun;21(2):301-13. PubMed PMID: 10907590.

33. Samarasinghe AE, Hoselton SA, Schuh JM. A comparison between intratracheal and inhalation delivery of Aspergillus fumigatus conidia in the development of fungal allergic asthma in C57BL/6 mice. Fungal biology. 2011 Jan;115(1):21-9. PubMed PMID: 21215951. Pubmed Central PMCID: 3053007.

34. Smith AM, Adler FR, Ribeiro RM, Gutenkunst RN, McAuley JL, McCullers JA, et al. Kinetics of coinfection with influenza A virus and Streptococcus pneumoniae. PLoS pathogens. 2013 Mar;9(3):e1003238. PubMed PMID: 23555251. Pubmed Central PMCID: 3605146.

35. Ghoneim HE, Thomas PG, McCullers JA. Depletion of alveolar macrophages during influenza infection facilitates bacterial superinfections. Journal of immunology. 2013 Aug 1;191(3):1250-9. PubMed PMID: 23804714.

36. McCullers JA. The co-pathogenesis of influenza viruses with bacteria in the lung. Nature reviews Microbiology. 2014 Apr;12(4):252-62. PubMed PMID: 24590244.

37. Young VB. The role of the microbiome in human health and disease: an introduction for clinicians. Bmj. 2017 Mar 15;356:j831. PubMed PMID: 28298355.

38. Busse WW, Gern JE. Asthma and infections: is the risk more profound than previously thought? The Journal of allergy and clinical immunology. 2014 Aug;134(2):260-1. PubMed PMID: 25087225.

39. Maccioni L, Weber S, Elgizouli M, Stoehlker AS, Geist I, Peter HH, et al. Obesity and risk of respiratory tract infections: results of an infection-diary based cohort study. BMC public health. 2018 Feb 20;18(1):271. PubMed PMID: 29458350. Pubmed Central PMCID: 5819164.

40. Falagas ME, Kompoti M. Obesity and infection. The Lancet Infectious diseases. 2006 Jul;6(7):438-46. PubMed PMID: 16790384.

41. Martin TR. The relationship between malnutrition and lung infections. Clinics in chest medicine. 1987 Sep;8(3):359-72. PubMed PMID: 3117482.

42. Hilliard KL, Allen E, Traber KE, Yamamoto K, Stauffer NM, Wasserman GA, et al. The Lung-Liver Axis: A Requirement for Maximal Innate Immunity and Hepatoprotection during Pneumonia. Am J Respir Cell Mol Biol. 2015 Sep;53(3):378-90. PubMed PMID: 25607543. Pubmed Central PMCID: PMC4566062. Epub 2015/01/22.

43. Carter R, Wolf J, van Opijnen T, Muller M, Obert C, Burnham C, et al. Genomic analyses of pneumococci from children with sickle cell disease expose host-specific bacterial adaptations and deficits in current interventions. Cell host & microbe. 2014 May 14;15(5):587-99. PubMed PMID: 24832453. Pubmed Central PMCID: 4066559.

44. Karlsson EA, Oguin TH, Meliopoulos V, Iverson A, Broadnax A, Yoon SW, et al. Vascular Permeability Drives Susceptibility to Influenza Infection in a Murine Model of Sickle Cell Disease. Sci Rep. 2017 Mar 3;7:43308. PubMed PMID: 28256526. Pubmed Central PMCID: PMC5335717. Epub 2017/03/04.

45. Rosch JW, Boyd AR, Hinojosa E, Pestina T, Hu Y, Persons DA, et al. Statins protect against fulminant pneumococcal infection and cytolysin toxicity in a mouse model of sickle cell disease. J Clin Invest. 2010 Feb;120(2):627-35. PubMed PMID: 20093777. Pubmed Central PMCID: PMC2810080. Epub 2010/01/23.

46. van Opijnen T, Camilli A. A fine scale phenotype-genotype virulence map of a bacterial pathogen. Genome Res. 2012 Dec;22(12):2541-51. PubMed PMID: 22826510. Pubmed Central PMCID: PMC3514683. Epub 2012/07/25.

47. Hava DL, Camilli A. Large-scale identification of serotype 4 Streptococcus pneumoniae virulence factors. Mol Microbiol. 2002 Sep;45(5):1389-406. PubMed PMID: 12207705. Pubmed Central PMCID: PMC2788772. Epub 2002/09/05.

48. Schrauwen EJ, de Graaf M, Herfst S, Rimmelzwaan GF, Osterhaus AD, Fouchier RA. Determinants of virulence of influenza A virus. Eur J Clin Microbiol Infect Dis. 2014 Apr;33(4):479-90. PubMed PMID: 24078062. Pubmed Central PMCID: PMC3969785. Epub 2013/10/01.

49. Schrauwen EJ, Fouchier RA. Host adaptation and transmission of influenza A viruses in mammals. Emerg Microbes Infect. 2014 Feb;3(2):e9. PubMed PMID: 26038511. Pubmed Central PMCID: PMC3944123. Epub 2014/02/01.

50. Tscherne DM, Garcia-Sastre A. Virulence determinants of pandemic influenza viruses. J Clin Invest. 2011 Jan;121(1):6-13. PubMed PMID: 21206092. Pubmed Central PMCID: PMC3007163. Epub 2011/01/06.

51. McCullers JA, Webster RG. A mouse model of dual infection with influenza virus and *Streptococcus pneumoniae*. International Congress Series. 2001;1219:601–7.

52. Morgan DJ, Casulli J, Chew C, Connolly E, Lui S, Brand OJ, et al. Innate Immune Cell Suppression and the Link With Secondary Lung Bacterial Pneumonia. Frontiers in immunology. 2018;9:2943. PubMed PMID: 30619303. Pubmed Central PMCID: 6302086.

53. Sun K, Salmon S, Yajjala VK, Bauer C, Metzger DW. Expression of suppressor of cytokine signaling 1 (SOCS1) impairs viral clearance and exacerbates lung injury during influenza infection. PLoS pathogens. 2014 Dec;10(12):e1004560. PubMed PMID: 25500584. Pubmed Central PMCID: 4263766.

54. Goulding J, Godlee A, Vekaria S, Hilty M, Snelgrove R, Hussell T. Lowering the threshold of lung innate immune cell activation alters susceptibility to secondary bacterial superinfection. The Journal of infectious diseases. 2011 Oct 1;204(7):1086-94. PubMed PMID: 21881124. Pubmed Central PMCID: 3164429.

55. Pettigrew MM, Gent JF, Kong Y, Wade M, Gansebom S, Bramley AM, et al. Association of sputum microbiota profiles with severity of community-acquired pneumonia in children. BMC infectious diseases. 2016 Jul 8;16:317. PubMed PMID: 27391033. Pubmed Central PMCID: PMC4939047. Epub 2016/07/09.

56. Kelly MS, Surette MG, Smieja M, Pernica JM, Rossi L, Luinstra K, et al. The Nasopharyngeal Microbiota of Children With Respiratory Infections in Botswana. Pediatr Infect Dis J. 2017 Sep;36(9):e211-e8. PubMed PMID: 28399056. Pubmed Central PMCID: PMC5555803. Epub 2017/04/12.

57. Sakwinska O, Bastic Schmid V, Berger B, Bruttin A, Keitel K, Lepage M, et al. Nasopharyngeal microbiota in healthy children and pneumonia patients. J Clin Microbiol. 2014 May;52(5):1590-4. PubMed PMID: 24599973. Pubmed Central PMCID: PMC3993659. Epub 2014/03/07.

58. Santee CA, Nagalingam NA, Faruqi AA, DeMuri GP, Gern JE, Wald ER, et al. Nasopharyngeal microbiota composition of children is related to the frequency of upper respiratory infection and acute sinusitis. Microbiome. 2016 Jul 1;4(1):34. PubMed PMID: 27364497. Pubmed Central PMCID: PMC4929776. Epub 2016/07/02.

59. Segal LN, Blaser MJ. A brave new world: the lung microbiota in an era of change. Annals of the American Thoracic Society. 2014 Jan;11 Suppl 1:S21-7. PubMed PMID: 24437400. Pubmed Central PMCID: 3972973.

60. Perez-Losada M, Authelet KJ, Hoptay CE, Kwak C, Crandall KA, Freishtat RJ. Pediatric asthma comprises different phenotypic clusters with unique nasal microbiotas. Microbiome. 2018 Oct 4;6(1):179. PubMed PMID: 30286807. Pubmed Central PMCID: PMC6172741. Epub 2018/10/06.

61. Beigelman A, Bacharier LB. Early-life respiratory infections and asthma development: role in disease pathogenesis and potential targets for disease prevention. Current opinion in allergy and clinical immunology. 2016 Apr;16(2):172-8. PubMed PMID: 26854761. Pubmed Central PMCID: 5089840.

62. von Mutius E, Vercelli D. Farm living: effects on childhood asthma and allergy. Nature reviews Immunology. 2010 Dec;10(12):861-8. PubMed PMID: 21060319.

63. von Mutius E. The microbial environment and its influence on asthma prevention in early life. The Journal of allergy and clinical immunology. 2016 Mar;137(3):680-9. PubMed PMID: 26806048.

64. Brooks C, Pearce N, Douwes J. The hygiene hypothesis in allergy and asthma: an update. Current opinion in allergy and clinical immunology. 2013 Feb;13(1):70-7. PubMed PMID: 23103806.

65. van Tilburg Bernardes E, Arrieta MC. Hygiene Hypothesis in Asthma Development: Is Hygiene to Blame? Archives of medical research. 2017 Nov;48(8):717-26. PubMed PMID: 29224909.

66. Risnes KR, Belanger K, Murk W, Bracken MB. Antibiotic exposure by 6 months and asthma and allergy at 6 years: Findings in a cohort of 1,401 US children. Am J Epidemiol. 2011 Feb 1;173(3):310-8. PubMed PMID: 21190986. Pubmed Central PMCID: PMC3105273. Epub 2010/12/31.

67. Murk W, Risnes KR, Bracken MB. Prenatal or early-life exposure to antibiotics and risk of childhood asthma: a systematic review. Pediatrics. 2011 Jun;127(6):1125-38. PubMed PMID: 21606151. Epub 2011/05/25.

68. Raymann K, Shaffer Z, Moran NA. Antibiotic exposure perturbs the gut microbiota and elevates mortality in honeybees. PLoS biology. 2017 Mar;15(3):e2001861. PubMed PMID: 28291793. Pubmed Central PMCID: 5349420.

69. Kim S, Covington A, Pamer EG. The intestinal microbiota: Antibiotics, colonization resistance, and enteric pathogens. Immunological reviews. 2017 Sep;279(1):90-105. PubMed PMID: 28856737. Pubmed Central PMCID: 6026851.

70. Lindenauer PK, Stefan MS, Feemster LC, Shieh MS, Carson SS, Au DH, et al. Use of Antibiotics Among Patients Hospitalized for Exacerbations of Asthma. JAMA Intern Med. 2016 Sep 1;176(9):1397-400. PubMed PMID: 27454705. Pubmed Central PMCID: PMC5515377. Epub 2016/07/28.

71. Francis KP, Yu J, Bellinger-Kawahara C, Joh D, Hawkinson MJ, Xiao G, et al. Visualizing pneumococcal infections in the lungs of live mice using bioluminescent Streptococcus pneumoniae transformed with a novel gram-positive lux transposon. Infection and immunity. 2001 May;69(5):3350-8. PubMed PMID: 11292758. Pubmed Central PMCID: 98294.

72. Hoselton SA, Samarasinghe AE, Seydel JM, Schuh JM. An inhalation model of airway allergic response to inhalation of environmental Aspergillus fumigatus conidia in sensitized BALB/c mice. Medical mycology. 2010 Dec;48(8):1056-65. PubMed PMID: 20482452. Pubmed Central PMCID: 3113699.

73. Andrews S. FastQC: A quality control tool for high throughput sequence data 2010. Available from: http://www.bioinformatics.babraham.ac.uk/projects/fastqc/

74. Krueger F. “Trim Galore.” A wrapper tool around Cutadapt and FastQC to consistently apply quality and adapter trimming to FastQ files 2015. Available from: http://www.bioinformatics.babraham.ac.uk/projects/trim_galore/.

75. Masella AP, Bartram AK, Truszkowski JM, Brown DG, Neufeld JD. PANDAseq: paired-end assembler for illumina sequences. BMC bioinformatics. 2012 Feb 14;13:31. PubMed PMID: 22333067. Pubmed Central PMCID: 3471323.

76. Caporaso JG, Kuczynski J, Stombaugh J, Bittinger K, Bushman FD, Costello EK, et al. QIIME allows analysis of high-throughput community sequencing data. Nature methods. 2010 May;7(5):335-6. PubMed PMID: 20383131. Pubmed Central PMCID: 3156573.

77. Edgar RC. Search and clustering orders of magnitude faster than BLAST. Bioinformatics. 2010 Oct 01;26(19):2460-1. PubMed PMID: 20709691.

78. DeSantis TZ, Hugenholtz P, Larsen N, Rojas M, Brodie EL, Keller K, et al. Greengenes, a chimera-checked 16S rRNA gene database and workbench compatible with ARB. Applied and environmental microbiology. 2006 Jul;72(7):5069-72. PubMed PMID: 16820507. Pubmed Central PMCID: 1489311.

79. McMurdie PJ, Holmes S. phyloseq: an R package for reproducible interactive analysis and graphics of microbiome census data. PLoS one. 2013;8(4):e61217. PubMed PMID: 23630581. Pubmed Central PMCID: 3632530.

80. Segata N, Izard J, Waldron L, Gevers D, Miropolsky L, Garrett WS, et al. Metagenomic biomarker discovery and explanation. Genome Biol. 2011 Jun 24;12(6):R60. PubMed PMID: 21702898. Pubmed Central PMCID: PMC3218848. Epub 2011/06/28.

